# Novel splicing and open reading frames revealed by long-read direct RNA sequencing of adenovirus transcripts

**DOI:** 10.1101/2019.12.13.876037

**Authors:** Alexander M. Price, Katharina E. Hayer, Daniel P. Depledge, Angus C. Wilson, Matthew D. Weitzman

## Abstract

Adenovirus is a common human pathogen that relies on host cell processes for production and processing of viral RNA. Although adenoviral promoters, splice junctions, and cleavage and polyadenylation sites have been characterized using low-throughput biochemical techniques or short read cDNA-based sequencing, these technologies do not fully capture the complexity of the adenoviral transcriptome. By combining Illumina short-read and nanopore long-read direct RNA sequencing approaches, we mapped transcription start sites and cleavage and polyadenylation sites across the adenovirus genome. The canonical viral early and late RNA cassettes were confirmed, but analysis of splice junctions within long RNA reads revealed an additional 20 novel viral transcripts. These RNAs include seven new splice junctions which lead to expression of canonical open reading frames (ORF), as well as 13 transcripts encoding for messages that potentially alter protein functions through truncations or the fusion of canonical ORFs. In addition, we also detect RNAs that bypass canonical cleavage sites and generate potential chimeric proteins by linking separate gene transcription units. Our work highlights how long-read sequencing technologies can reveal further complexity within viral transcriptomes.

## Introduction

Adenoviruses (AdV) are common viral pathogens across multiple species with distinct tissue tropisms including gut, eye, and lung [1]. Among the human adenoviruses, serotypes 2 (Ad2) and 5 (Ad5) from subgroup C are the most prevalent within the population, and they cause benign to severe respiratory infections [2]. These two serotypes are highly homologous, sharing 94.7% nucleotide identity between their genomes and 69.2-100% amino acid identity amongst conserved open reading frames (ORFs) [3,4]. AdVs readily infect most transformed human cell lines and have proven a valuable tool that has led to seminal discoveries in molecular biology for many decades [5]. RNA splicing was discovered by the analysis of adenovirus encoded RNAs [6,7], as well as other important findings in messenger RNA capping and polyadenylation [8,9]. It is now understood that essentially all AdV mRNAs are capped, spliced, polyadenylated, and exported from the nucleus using host cell machinery [10].

AdV are capable of infecting non-dividing cells and reprogramming cellular processes for productive viral infection. This rewiring involves a highly regulated cascade of viral gene expression over various kinetic classes [5]. The first viral gene to be expressed after infection is E1A, a multi-functional transcription factor that activates downstream viral transcription, liberates E2F from RB proteins, as well as alters host transcriptional responses to the virus [11–14]. While all E1A molecules have identical 5’ and 3’ nucleotide sequences, splicing of differently sized internal introns allows for the production of discrete proteins that lack specific functional domains conserved across serotypes [15]. Early after infection, E1A is expressed mainly as large and small isoforms, but later in infection alternative splicing leads to the production of a 9 Svedberg E1A isoform (E1A-9s) as well as low abundance doubly-spliced E1A-11s and E1A-10s. The second viral gene to be activated is E1B, consisting of predominantly two spliced isoforms producing 19-kilodalton and 55-kilodalton proteins, with two less abundant isoforms generating putative ORFs of 156 and 93 residues [16]. While E1B-19K acts to block cellular apoptosis [17], E1B-55K is another multifunctional protein that can cooperate with E1A to alter cellular gene expression downstream of p53 as well as form the targeting component of a viral ubiquitin ligase [18–23]. The remaining early transcription units are all transcriptionally activated by E1A and encode for products of related function. The E2 region on the reverse strand of the AdV genome has both an early and a late promoter, as well as two distinct polyadenylation sites, leading to upstream E2A and downstream E2B transcripts [24]. E2A encodes for the viral DNA-binding protein (DBP), while alternative splicing to E2B encodes for the protein-priming terminal protein (pTP) as well as the AdV DNA polymerase (AdPol) [25–27]. The E3 region encoded on the top strand also has two polyadenylation sites leading to E3A and E3B transcription units, and these gene products are primarily involved in modulating the host innate immune system [28–30]. Like E1A, the E4 region on the reverse strand has identical 5’ and 3’ regions, and encodes up to six ORFs by removal of a first intron of varying length. E4 region transcripts encode for multifunctional proteins that are involved in regulation of transcription, splicing, and translation of viral RNAs, as well as antagonizing intrinsic cellular defenses [31–33]. Additionally, AdV encodes two Pol III-derived virus associated (VA) RNAs involved in the inactivation of Protein Kinase acting on RNA (PKR) [34,35]. Ultimately, the concerted efforts of the AdV early proteins lead to a cellular state that allows for the replication and amplification of the viral DNA genome [36].

Prior to viral DNA replication, the AdV Major Late Promoter (MLP) is thought to be largely silent with small amounts of RNA being made that terminate at the immediately downstream (L1) polyadenylation site [37]. At this time, so-called intermediate genes pIX and IVa2 begin to be expressed from promoters within the E1B cassette and antisense to the MLP. Both pIX and IVa2 co-terminate at polyadenylation sites within the early genes they overlap with (E1B and E2B, respectively) and are involved in late gene transcription and packaging [38,39]. Only after viral DNA replication has occurred does the MLP fully activate, supporting the hypothesis that that active replication *in cis* is a prerequisite for full viral late gene expression [40–42]. The Major Late Transcriptional Unit (MLTU) begins with a series of three constitutive exons spliced together to form the tripartite leader, before downstream splicing to late cassettes defined by one of five alternative polyadenylation sites (termed L1-L5) [37]. Splicing within the tripartite leader to the so-called “i” exon leads to a putative ORF upstream of subsequent late gene splicing events and destabilizes these RNA molecules [43,44]. An additional intermediate promoter has been reported within the L4 region that allows for the early expression of L4-22K and L4-33K proteins important for the splicing of other late genes [45,46]. The MLTU encodes for primarily structural capsid components or proteins involved with packaging of new virions, and their expression ultimately leads to the death of the host cell. Recently, a novel late gene, UXP, was discovered on the reverse strand of the genome [47,48]. The UXP promoter is located between E4 and E2 on the reverse strand of the genome, and splices downstream to the exons within the E2A region to continue translation of an ORF in an alternate reading frame to that of DBP. This exciting finding suggests that our knowledge of AdV transcripts is incomplete, especially within the complex MLTU region.

The Ad5 genome was fully sequenced in 1991 using Sanger sequencing of viral genome fragments inserted into plasmid DNA and amplified in bacteria [3]. This genome sequence was then annotated in 2003 based on homology to similar serotypes of AdV [4]. As such, the current reference annotation for Ad5 available on the National Center for Biotechnology Information (<u>AC_000008</u>) is incomplete, and lacks critical information such as transcription start sites (TSS), cleavage and polyadenylation sites (CPAS), and the resulting 5’ and 3’ untranslated regions (UTR) that the aforementioned information dictates. In recent years, new technologies have allowed for high-throughput investigation of gene expression utilizing various techniques. The effect of AdV infection on host gene expression has been shown for Ad5 by microarray analysis [49,50], as well as for Ad2 by Illumina-based short-read sequencing [51,52]. Analyses of both single-end and paired-end short-read RNA-seq data from cells infected with Ad2 revealed both temporal viral gene expression and high-depth splicing information and identified both previously confirmed and novel RNA splice site junctions [53]. In addition. temporal analysis of Ad5 viral gene expression was performed using digital PCR to determine expression kinetics of a subset of known viral genes [54]. Lastly, the late RNA tripartite leader splicing was analyzed by short-read sequencing across a number of human AdV serotypes [43]. To date, no group has performed a comprehensive analysis of the RNAs generated during Ad5 infection. Furthermore, even though the quality and depth of current short-read sequencing technologies is high, the complex nature of many viral transcriptomes precludes the unambiguous mapping of these short reads to any one particular RNA isoform due to extreme gene density and overlapping transcriptional units [55,56]. In this regard, the ability of long-read RNA sequencing to map full-length transcripts has the potential to revolutionize detection of divergent isoforms and multiply spliced RNA at the single-molecule level [57–59].

In this study, we have re-annotated the Ad5 genome and transcriptome using a combination of short-read and long-read RNA sequencing technologies. The high read depth and accuracy of base-calling achieved by Illumina-based short-read sequencing allowed for both the detection of single nucleotide polymorphisms within transcriptionally active regions of the viral DNA genome, as well as error-correction of the inherently noisier base-calling of Nanopore-based long-read direct RNA sequencing (dRNA-seq). dRNA-seq enabled the detection of full-length RNA transcripts and the assignment of TSS and CPAS transcriptome-wide. Furthermore, by combining highly accurate splice site junctions from short-read sequencing and full-length isoform context from long-read sequencing, we were able to reevaluate the splicing complexity of AdV transcriptional units. Using this integrated approach, we have discovered 20 additional viral polyadenylated RNAs for a total of 75 unique mRNAs produced by Ad5. Of these novel isoforms, seven RNAs encode for a canonical ORF with changes in upstream or downstream splicing. The remaining 13 encode new ORFs or alter existing ORFs by internal truncations or in-frame fusion of genes from separate transcription units. Taken together, our data reveal additional transcriptional complexity of AdV and highlight the necessity of revisiting transcriptome annotations following the emergence of appropriate new technologies.

## Results

### RNA-seq reveals high-confidence SNPs within the Ad5 genome

Illumina-based RNA sequencing (RNA-seq) relies on the fractionation of RNA molecules before reverse transcription into complementary DNA, and therefore loses information such as RNA modifications and the context of splice junctions within full length molecules. However, the accuracy of each individual base call is very high [60]. Using bcftools, a common variant-calling algorithm designed to assess allele-specific variation within RNA-seq, we were able to detect single nucleotide polymorphisms (SNPs) within the RNA transcriptome that likely emerge from mutation within the DNA genome [61,62]. While RNA modifications such as inosine can be read as SNPs during the process of reverse transcription, these events should not approach the near 100% read depth stringency we required among our three biological replicates to call a conserved variant [63]. While this technique is only applicable for the actively transcribed region of the genome, nearly every nucleotide of the gene-dense AdV genome is transcribed at a sufficient level for this strategy to provide meaningful data.

In total, we discovered 24 SNPs and no insertions or deletions in the Ad5 genome when compared to the original annotation (**Figure 1**). Of these mutations, exactly half (12) are not predicted to change amino acid coding capacity, with two SNPs occurring within untranslated regions of viral RNA and the remaining ten leading to synonymous amino acid codons within all reading frames annotated to be protein producing. The remaining 12 mutations are predicted to lead to coding sequence variations at the amino acid level, with all examples being missense mutations and no evidence of premature stop codons. Importantly, none of the mutations discovered generated novel RNA splice sites. These data demonstrate the ability to call mutations within the DNA genomes of viruses using solely high-depth RNA sequencing data. Furthermore, detecting only 24 SNPs out of 35,938 nucleotides highlights the overall genomic stability of AdV.

**Figure 1.**
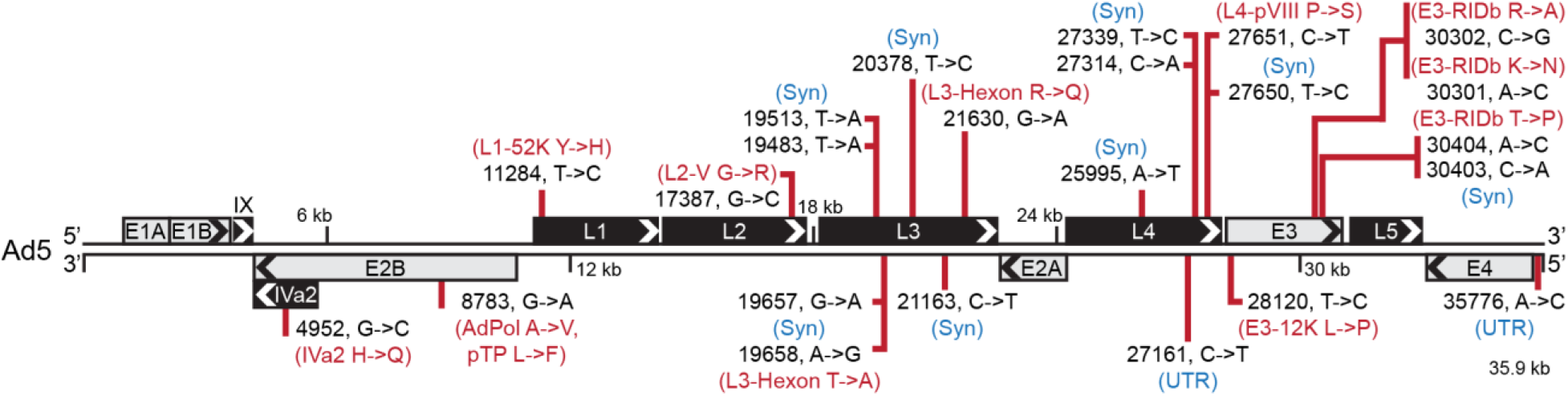
RNA-seq reveals high-confidence SNPs within the Ad5 genome. The 35,938 base pair linear genome of Ad5 is displayed in the traditional left to right format. Major transcriptional units are shown as boxes above or below the genome with arrowheads denoting the orientation of the open reading frames (ORFs) encoded within. Grey boxes denote early gene transcriptional units while black boxes denote late genes. Bcftools was used to analyze short-read RNA seq data to predict single nucleotide polymorphisms (SNPs) and insertions/deletions (InDels) that approach 100% of the RNA reads when compared to the reference Ad5 genome (AC_000008). In total, 23 such SNPs were discovered and their position within the genome is highlighted by a red vertical line. For each SNP, the nucleotide position as well as the top strand reference base and corrected base are shown in black text (nucleotide position, reference base -> corrected base). If indicated SNPs fell within untranslated regions (UTR), or did not change the encoded amino acid of any annotated reading frame potentially impacted by the SNP, these were marked with blue text denoting either UTR or Syn (synonymous mutation), respectively. For any SNP that led to an amino acid change within an annotated ORF, these ORFs as well as the identity of the reference amino acid and corrected amino acid are highlighted in red.

### Combined short-read and long-read sequencing showcases adenovirus transcriptome complexity

To compare short-read Illumina sequencing and long-read nanopore sequencing directly, A549 cells were infected with Ad5 for 24 hours and total RNA was harvested in biological triplicate. Fractions of these three samples were prepared into standard strand specific Illumina RNA-seq libraries using the polyadenylated mRNA fraction. The same RNA samples were then poly(A) purified before submitting to direct RNA sequencing (dRNA-seq) on an Oxford Nanopore Technologies MinION MkIb platform [64]. Resulting sequence reads were aligned to the Ad5 reference genome using either GSNAP for short-reads [65], or MiniMap2 for long-reads [66]. Overall sequencing depth for both forward and reverse reads are shown in **Figure 2**. While Illumina sequencing provided on average three times the read depth when compared to dRNA sequencing, the overall coverage plots were similar.

**Figure 2.**
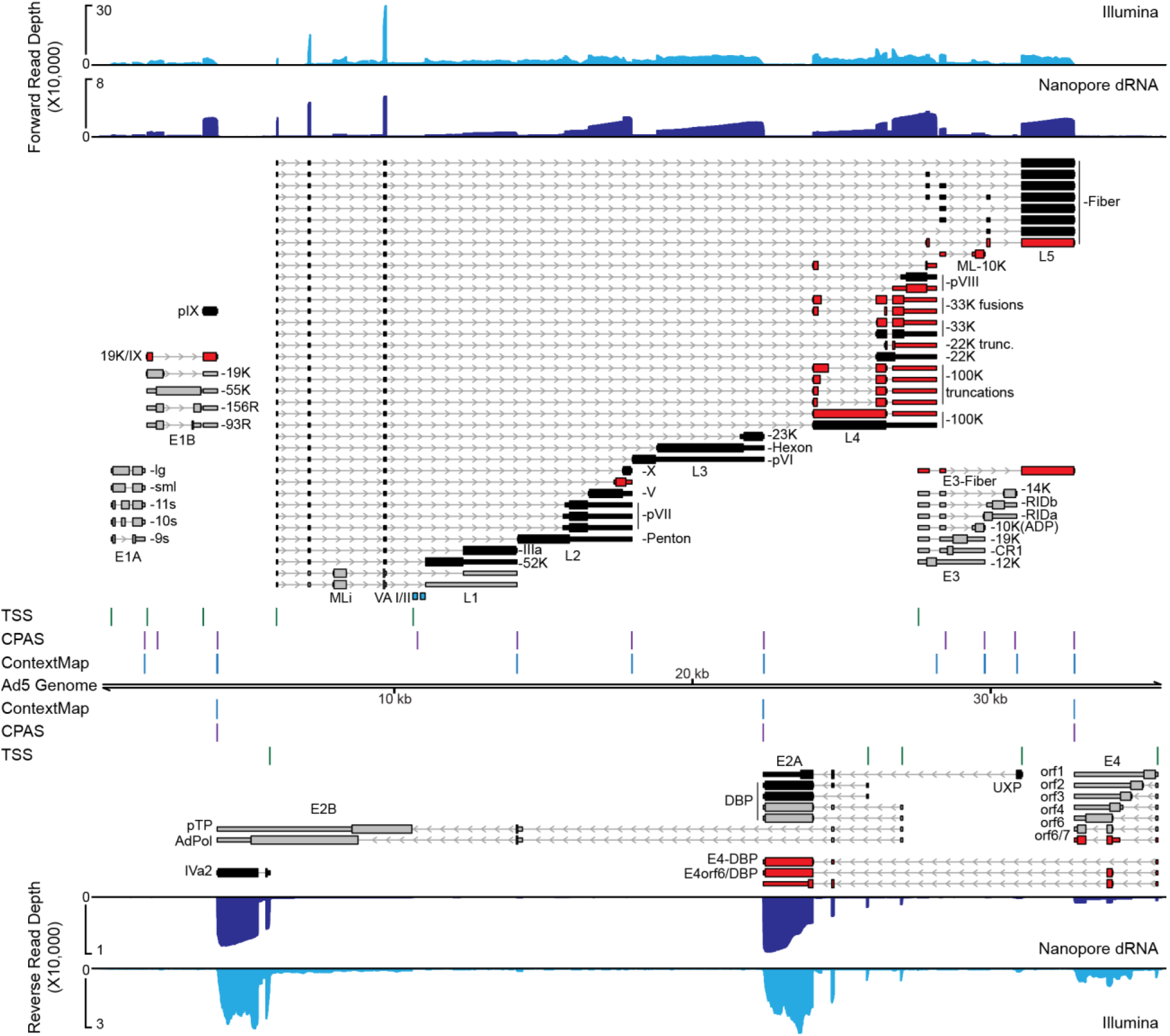
Combined short-read and long-read sequencing showcases adenovirus transcriptome complexity. A549 cells were infected with Ad5 for 24 hours before RNA was extracted and subjected to both short-read and long-read sequencing. Sequence coverage provided by short-read stranded RNA-seq (Illumina, light blue), as well as nanopore long-read direct RNA-seq (Nanopore dRNA, dark blue), is shown along the Ad5 genome. For both tracks, reads aligning to the forward strand are plotted above the genome, while reads aligning to the reverse strand are shown below. For dRNA-seq datasets, reads can be reduced to their 5’ and 3’ ends and peak-calling applied to predict individual transcription start sites (TSS, green vertical lines) or cleavage and polyadenylation sites (CPAS, magenta vertical lines), respectively. Similarly, the ContextMap algorithm can predict, albeit at lower sensitivity, CPAS sites from poly(A) containing fragments within Illumina RNA-seq data (ContextMap, light blue vertical lines). Individual RNA transcripts are shown above and below the genome, thin bars denote 5’ and 3’ untranslated regions (UTR), thick bars denote open reading frames (ORFs), and thin lines with arrowheads denote both introns and orientation of transcription. Previously characterized early genes are denoted in grey, while previously characterized late genes are denoted in black. RNA isoforms discovered in this study are highlighted in red. Names of transcriptional units are shown under each cluster of transcripts, while the name of the protein derived from the respective ORF is listed after each transcript. The position of Pol III-derived noncoding RNAs virus associated VA-I and VA-II are highlighted in teal boxes.

dRNA-seq is performed in the 3’ -> 5’ direction and thus allows precise mapping of the 3’ ends of transcripts at which poly(A) tails are added (cleavage and polyadenylation site, CPAS) [64]. Where the quality of input RNA is high, a variable proportion of sequence reads extend all the way to their transcription start site (TSS). By collapsing sequences reads to their 5’ and 3’ ends, we were able to implement a peak-calling approach to predict TSS and CPAS [57], and map their positions along both the forward and reverse strand of the viral genome (**Figure 2**). In addition, ContextMap2 [67] was used to mine Illumina RNA-seq data for short read sequences containing poly(A) stretches that could be aligned against the viral genome for an orthogonal method of CPAS detection (**Figure 2**). Mapping the TSS on the forward strand revealed the locations of the E1A, E1B, pIX, MLP, and E3 promoters, while the reverse strand revealed the E4, UXP, E2-early, E2-late, and IVa2 promoters. We did not detect any transcripts starting internal to L4 at the proposed L4 promoter [45]. When mapping CPAS loci, we saw great concordance between the dRNA-seq and ContextMap2 performed on short-read sequences. On the forward strand we were able to detect previously mapped CPAS events at the E1A, E1B/pIX, E3A, E3B, and individual L1 through L5 sites. On the reverse strand we detected CPAS at the E4, UXP/DBP, and E2B/IVa2 locations. In addition, we also detected TSS and CPAS around the RNA pol III-derived VA RNA I (**Figure 2**). While pol III transcripts are generally not polyadenylated, and thus would not be captured by our nanopore sequencing approach, it was previously reported that low levels of polyadenylation can occur on these transcripts [68]. Given the high abundance of AdV VA RNAs (up to 10^8^ copies per cell during late infection), it remains likely that low level VA RNA polyadenylation events are occurring [35].

To generate accurate splicing maps of AdV transcripts we combined the sensitivity of short-read sequencing to identify RNA junctions and then placed them in the context of full-length RNA isoforms using dRNA sequencing. Due to the spurious nature of low-level AdV splicing events [53], we set abundance thresholds for the highly abundant viral late transcripts of 500 reads for short-read junctions, and at least ten events detected in the long-read sequencing when collapsed by FLAIR [69]. Using this method, we readily detected other recently discovered viral isoforms, such as multiple splice sites preceding the pVII ORF [53], the so-called X, Y, and Z leaders embedded in E3 and preceding L5-Fiber [53,70], and the newly described UXP [47,48]. Using full-length RNAs, we were able to detect novel splice sites producing canonical ORFs that only differ in UTRs for L4-100K, L4-33K, L4-pVIII, and E4orf6/7. In addition, we discovered canonical ORF isoforms embedded within transcripts generated from non-canonical promoters, such as Fiber driven by the E3 promoter, E3-10K driven by the Major Late Promoter, and DBP driven by the E4 promoter. Within transcriptional units, we discovered the presence of internal splice sites leading to in-frame truncations of existing ORFs, such as L4-22K and four distinct isoforms of truncated L4-100K. We also discovered splicing events predicted to lead to in-frame fusion events within transcriptional units, such as fusions between N-terminal fragments of L4-100K and L4-33K or L4-pVIII or the X-Z-Fiber ORF. Furthermore, gene fusion events were observed that join disparate transcriptional units, such as an N-terminal fragment of E1B-19K and pIX (19K/IX) or E4orf6 and DBP (E4orf6/DBP). Lastly, we detected a splice site leading to a novel ORF of predicted 13 kilodaltons (L2-Unk13K) between the splice sites for L2-V and L2-pX. This novel L2 splice site was conserved in Ad2 [53]. Overall, we discovered 20 new isoforms for a total of 75 expressed RNA isoforms during Ad5 infection. Of these, many potentially exciting fusions and truncations of existing ORFs remain to be explored.

### Direct RNA Sequencing unambiguously distinguishes early and late transcription

We next determined if we could provide unambiguous detection of viral transcripts over a time-course of infection that recapitulated early and late viral kinetics. By aligning long reads to the fully re-annotated viral transcriptome (as opposed to the viral genome), and only counting the reads that could be unambiguously assigned to a single transcript, we were able to detect all of the canonical and newly discovered transcripts (**Figure 3**). At 12 hours post infection (hpi) the majority of viral transcripts detected were early RNAs, particularly E1A-large and E1A-small, E1B-19K and E1B-55K, early promoter DBP, and E4orf3 (**Figure 3A**). However, at this time point we still detected low-level viral late transcripts that progressed beyond the L1 polyadenylation site, corroborating recent work [54]. At 24 hpi, however, viral gene expression shifted to be dominated by late gene expression, as well as early transcripts derived from the E1B locus (**Figure 3B**). At late times post infection we also saw the E3-Fiber transcript, 19K/IX, and E4-DBP transcripts increase dramatically, potentially implicating these messages as novel late transcripts, with expression as abundant as the recently described late UXP transcript [47,48]. Furthermore, while all permutations of the X, Y, and Z leaders preceding Fiber were previously detected by short-read sequencing these could not be phased to full-length transcript isoforms [53,70]. Our full-length RNA data indicate that all Fiber transcripts can be detected, but MLP-Fiber and Y-Fiber are the most abundant, followed by XY-Fiber, and then all other isoforms. While the previous lack of detection of some of these novel transcripts can be explained by low overall abundance (e.g., L2-Unk13K, L4-100K/VIII) many of the L4-100K truncations and L4-33K fusions are expressed at levels higher than that of the bona fide late transcript UXP. These data demonstrate that the newly discovered viral transcripts can be reproducibly detected over a time-course of infection with Ad5, as well as display differential expression based on the stage of infection.

**Figure 3.**
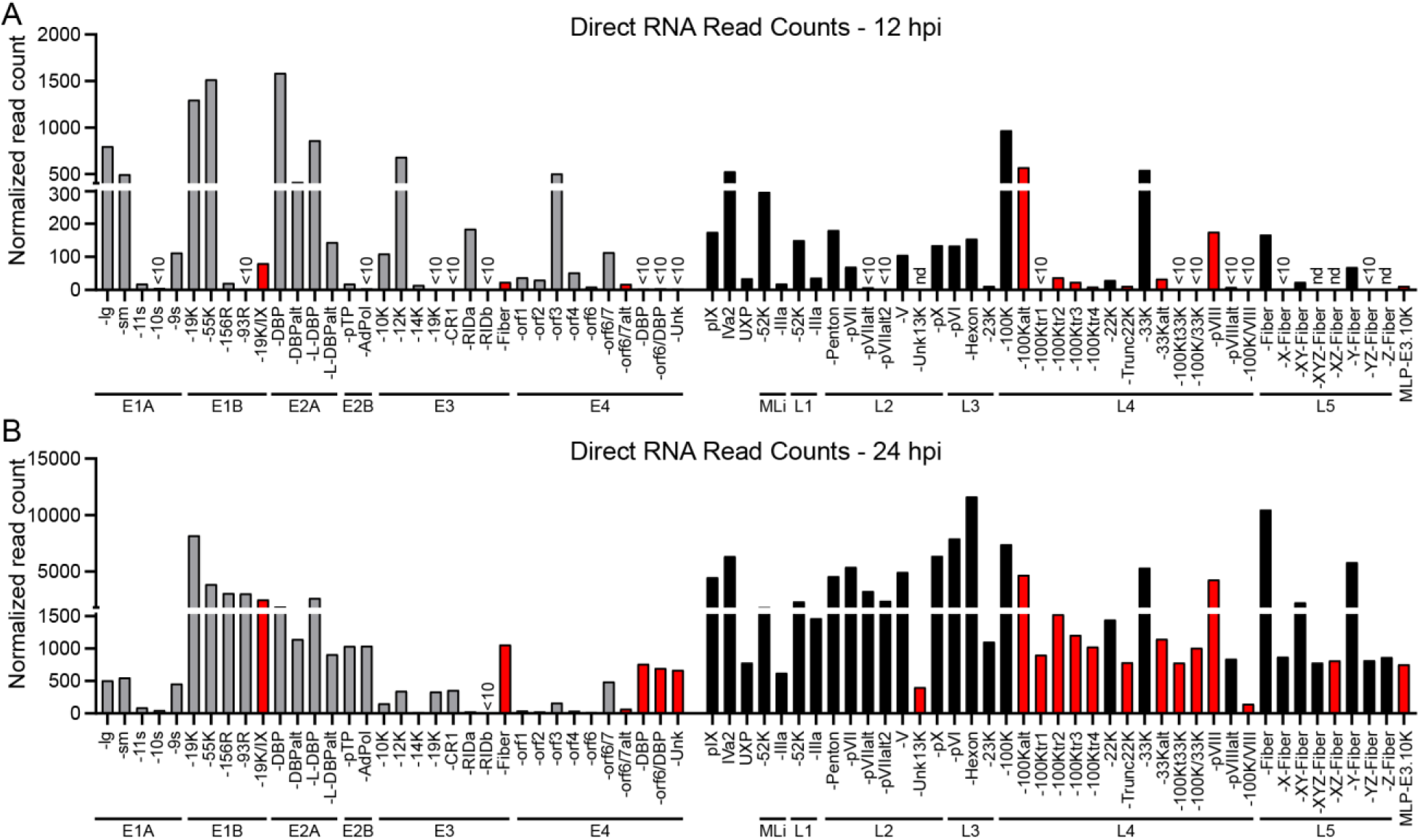
Direct RNA Sequencing (dRNA-seq) unambiguously distinguishes early and late transcription. **(A)** dRNA-seq was performed on polyadenylated RNA from Ad5-infected A549 cells extracted at 12 hours post infection (hpi). Sequence reads were aligned to the re-annotated transcriptome and filtered to retain only unambiguous primary alignments. Normalized read count indicates the number of RNAs for a particular transcript once normalized to the total number of mappable reads (human plus adenovirus) for the entire sequencing reaction. For all panels, grey bars indicate early genes, black bars indicate late genes, and red bars indicate novel isoforms discovered in this study. Particular transcripts are highlighted if there were less than 10 counts of a particular isoform detected (<10), or if the RNA was undetectable at that time point (nd). **(B)** Same as in Panel A, but with RNA harvested at 24 hpi.

## Discussion

DNA viruses encode large amounts of information in compact genomes through alternative splicing, overlapping transcripts, and transcription from both strands of the genome. The complexity of the adenovirus transcriptome has not been fully explored using modern high-throughput technologies. Here we integrate short-read cDNA sequencing and long-read direct RNA sequencing to re-annotate both the Ad5 DNA genome and RNA transcriptome. Using high quality and high depth short-read sequencing, we were able to detect SNPs within the transcribed regions of the genome approaching 100% penetrance, indicating that these sites were likely present in the genome and not due to RNA editing or modifications. We recapitulated the known TSS and CPAS sites throughout the Ad5 genome, and annotated novel splicing events within the viral transcriptome. Of these 20 novel RNAs, 13 are likely to encode for altered ORFs including multiple fusion transcripts that span transcriptional units thought previously to be distinct. Overall, we have provided a more complete annotation of a complex viral transcriptome that highlights potentially new gene products for future study.

Using RNA-seq data to call SNPs in viral DNA genomes is compelling since high quality short-read sequencing data sets already exist for many DNA viruses [71–75]. While half of the SNPs we called were synonymous or in non-coding regions, missense mutations have the potential to change the coding sequence of protein amino acids in meaningful ways. In addition, while SNPs are often tolerated during alignment of RNA-seq data, annotation of the correct primary amino acid sequence is critical for downstream analysis of mass spectrometry data [76]. While the SNPs we detected might be bona fide mutations that have arisen during passage in cell culture, it is also possible that the original reference sequence contains errors introduced by the sequencing technologies employed at the time [3,4]. It will be critical to directly sequence the DNA genomes of Ad5 isolates from multiple laboratories to test this hypothesis.

Previous studies of AdV transcription detected numerous splice sites beyond those employed by known isoforms. However, the constraints of short-read sequencing precluded proper assembly of these sites into full-length transcripts [43,53,70]. Furthermore, targeted expression analysis over a time-course of infection was limited to already known transcripts [54]. Using direct RNA sequencing we have been able to confirm that these RNAs exist (e.g., the various x, y, and z leaders preceding some molecules of Fiber transcripts), as well as show regulated expression over a time-course of infection. We have also added ORF predictions to previously detected splice sites, such as L2-Unk13K, X-Z-Fiber, and the pVIII ORF derived from splicing directly from the tripartite leader to the L4-33K splice acceptor. While this last site was previously predicted to lead to the expression of a small 42 amino acid ORF [53], we propose that this transcript instead primarily encodes for pVIII with a small upstream ORF, as it is over five times as abundant as the canonical pVIII spliced RNA. It should be noted that we did not detect the presence of a putative L4 intermediate promoter TSS at either 12 or 24 hpi [45,77,78]. One hypothesis is that the sequence detected in L4-100K that is necessary for early expression of L4-22K and L4-33K might instead encode for a *cis*-regulatory element that mediates the early accumulation of these two products produced from the major late promoter.

Of the novel transcripts we have so far detected, all of them appear to display delayed late kinetics during infection. Of particular interest is the transcript encoding for a putative fusion event between the E4 transcriptional unit and the E2 transcriptional unit. This transcript, E4orf6/DBP, would have to skip the canonical E4 CPAS for the pre-mRNA to progress downstream to DBP for splicing. The three transcripts displaying this pattern, including E4-promoter driven DBP and frameshifted E4-Unk, are all much more abundant during the late phase of infection even though canonical E4 transcripts are expressed early. It will be very interesting to see if differential polyadenylation is regulated during the life cycle of the virus, as has been previously reported for herpes simplex virus [57,74,79]. Importantly, future research should identify whether the known functions of existing viral ORFs can be explained, at least in part, by the presence of these novel isoforms.

## Methods

### Cell Culture

A549 cells (ATCC CCL-185) were obtained from American Type Culture Collection (ATCC) and cultured at 37 °C and 5% CO2. Cells were maintained in Ham’s F-12K medium (Gibco, 21127-022) supplemented with 10% v/v FBS (VWR, 89510-186) and 1% v/v Pen/Strep (100 U/ml of penicillin, 100 μg/ml of streptomycin, Gibco, 15140-122). All cell lines tested negative for mycoplasma infection and were routinely tested afterwards using the LookOut Mycoplasma PCR Detection Kit (Sigma-Aldrich).

### Viral infections

Adenovirus serotype 5 (Ad5) was originally purchased from ATCC. All viruses were expanded on HEK293 cells, purified using two sequential rounds of ultracentrifugation in CsCl gradients, and stored in 40% v/v glycerol at −20 °C (short term) or −80 °C (long term). Viral stock titer was determined on HEK293 cells by plaque assay, and all subsequent infections were performed at a multiplicity of infection (MOI) of 10 PFU/cell. Cells were infected at 80-90% confluent monolayers by incubation with diluted virus in a minimal volume of low serum (2%) F-12K for two hours. After infection viral inoculum was removed by vacuum and full serum growth media was replaced for the duration of the experiment.

### RNA Isolation

Total RNA was isolated from cells by either TRIzol extraction (Thermo Fisher) or RNeasy Micro kit (Qiagen), following manufacturer protocols. RNA was treated with RNase-free DNase I (Qiagen), either on-column or after ethanol precipitation. To test quality, RNA was converted to complementary DNA (cDNA) using 1 μg of input RNA in the High Capcity RNA-to-cDNA kit (Thermo Fisher). Quantitative PCR was performed using the standard protocol for SYBR Green reagents (Thermo Fisher) in a QuantStudio 7 Flex Real-Time PCR System (Applied Biosystems).

### Illumina Sequencing and Mapping

Total RNA from three biological replicates of Control knockdown or three biological replicates of METTL3-knockdown A549 cells infected with Ad5 for 24 hours were sent to Genewiz for preparation into strand-specific RNA-Seq libraries. Libraries were then run spread over three lanes of an Illumina HiSeq 2500 using a 150bp paired-end protocol. Raw reads were mapped to the GRCh37/hg19 genome assembly and the Ad5 genome using the RNA-seq aligner GSNAP [65] (version 2019-09-12). The algorithm was given known human gene models provided by GENCODE (release_27_hg19) to achieve higher mapping accuracy. We used R package ggplot2 for visualization. Downstream analysis and visualization was done using deepTools2 [80]. Splice junctions were extracted using regtools [81] and visualized in Integrative Genomics Viewer [82].

### Variant Calling

Illumina RNA-seq reads were aligned to the Ad5 genome obtained from NCBI (https://www.ncbi.nlm.nih.gov/nuccore/AC_000008) using GSNAP [65]. To identify variants such as single nucleotide polymorphisms (SNPs) and insertions/deletions (InDels), we combined mpileup and call from the bcftools (v1.9) package [61,62]. Here we used the following flags “-- redo-BAQ --min-BQ 30 --per-sample-mF” and “--multiallelic-caller --variants-only” respectively. Finally, we only considered variants if they were called significantly in all 3 replicates. We only observed SNPs but no InDels.

### Direct RNA Sequencing on nanopore arrays

Direct RNA sequencing libraries were generated from 800-900 ng of poly(A) RNA, isolated using the Dynabeads™ mRNA Purification Kit (Invitrogen, 61006). Isolated poly(A) RNA was subsequently spiked with 0.3 μl of a synthetic Enolase 2 (ENO2) calibration RNA (Oxford Nanopore Technologies Ltd.) and prepared for sequencing using standard protocol steps previously described [57,64]. Sequencing was carried out on a MinION MkIb with R9.4.1 (rev D) flow cells (Oxford Nanopore Technologies Ltd.) for 20 hours and generated 550,000-770,000 sequence reads per dataset. Raw fast5 datasets were basecalled using Guppy v3.2.2 (-f FLO-MIN106 -k SQK-RNA002) and subsequently aligned against the adenovirus Ad5 reference genome (AC_000008.1) using MiniMap225 (-ax splice -k14 -uf --secondary=no), a splice aware aligner [66]. Resulting SAM files were parsed using SAMtools v1.3 [83].

### Defining TSS and CPAS

Transcription start sites (TSS) as well as RNA cleavage and polyadenylation sites were identified as follows. Sorted BAM files containing sequence reads aligned to the Ad5 genome were parsed to BED12 files using BEDtools [84], separated by strand, truncated to their 5’ and 3’ termini, and output as BED6 files. Peak regions denoting TSS and CPAS were identified using the HOMER [85] findpeaks module (-o auto -style tss) using a --localSize of 100 and 500 and --size of 15 and 50 for TSS and CPAS, respectively. TSS peaks were compared against Illumina annotated splice sites to identify and remove peak artefacts derived from local alignment errors around splice junctions. To predict CPAS sites on the viral genome, we also used the RNA-seq aligner ContextMap2 (version 2.7.9) [67] which has poly(A) read mapping implemented on our short-read data. To run this tool, we used the following optional flags “-aligner_name bowtie --polyA --strandspecific”. Due to the previously reported errors when using ContextMap2 at very high read depth, we chose to randomly subsample 10 million, 20 million and 30 million and run the tool on each of the subsets. We only report poly(A) sites if they were called in all three replicates and in at least two of the subsample groups.

### Splice junction correction and sequence read collapsing

Illumina-assisted correction of splice junctions in direct RNA-Seq data was performed using FLAIR v1.3 [69] in a stranded manner. Briefly, Illumina reads aligning to the Ad5 genome were split according to orientation and mapping strand [-f83 & -f163 (forward) and -f99 & -f147 (reverse)] and used to produce strand-specific junction files that were filtered to remove junctions supported by less than 100 Illumina reads. Direct RNA-Seq reads were similarly aligned to the Ad5 genome and separated according to orientation [-F4095 (forward) and -f16 (reverse)] prior to correction using the FLAIR correct module (default parameters). Resulting BED12 files were parsed to extend the termini of each individual sequence read to the nearest TSS and CPAS with BlockStarts and BlockSizes (BED12 cols 11 & 12) corrected to reflect this. BED12 files were subsequently collapsed by identifying all reads sharing the same BlockStarts and BlockSizes and reducing these to a single representative. Resulting data were visualized along with the raw read data using IGV [82] and low abundance isoforms (supported by less than 500 junctional reads or 10 full-length reads from Illumina or nanopore data, respectively) removed prior to producing the final annotation.

### Isoform counting

Using our new Ad5 annotation, we generated a transcriptome database by parsing our GFF3 file to a BED12 file using the *gff3ToGenePred* and *genePredtoBED* functions within UCSCutils (https://github.com/itsvenu/UCSC-Utils-Download) and subsequently extracting a fasta sequence for each transcript isoform using the *getfasta* function within BEDtools [84]. Direct RNA-Seq reads were then aligned against the transcriptome database using parameters optimized for transcriptome-level alignment (minimap2 -ax map-ont -p 0.99). Isoform counts were generated by filtering only for primary alignments (SAM flag 0) with a mapping quality (MapQ) > 0.

## Acknowledgments

We thank members of the Weitzman and Mohr/Wilson Labs for insightful discussions and input. This work was supported through NIH grants R21-AI130618 and R21-AI147163 (ACW), and R01-AI145266, R01-AI121321, and R01-CA097093 (MDW). Additional support came from the NCI T32 Training Grant in Tumor Virology T32-CA115299 (AMP) and Individual National Research Service Award F32-AI138432 (AMP). We extend special thanks to Ian Mohr (New York University School of Medicine) for support of DPD in part through National Institutes of Health (NIH) grants R01-AI073898 and R01-GM056927.

## Data Availability

Basecalled fast5 (Nanopore) and fastq (Illumina) datasets generated as part of this study can be downloaded from the European Nucleotide Archive (ENA) under the following study accession: PRJEB35667. The authors declare that all other data supporting the findings of this study are available within the article and its Supplementary Information files, or are available from the authors upon request. The newly generated genome and transcriptome annotation can be found at https://github.com/dandepledge/Ad5-annotation.

## Author Contributions

A.M.P. and M.D.W. conceived of the project and designed the experiments; D.P.D. and A.C.W. provided additional input into study design; A.M.P. performed the experiments and Illumina sequencing; D.P.D. performed the nanopore sequencing; K.E.H. and D.P.D. performed computational analyses; A.M.P. and D.P.D analyzed all additional data; A.M.P. and M.D.W. wrote the manuscript; All authors read, edited, and approved the final paper.

## References

1. Arnold J. Berk. Adenoviridae. 6th ed. In: David M. Knipe, Peter M. Howley, editors. Fields Virology. 6th ed. Philadelphia: Wolters Kluwer Health/Lippincott Williams & Wilkins; 2013. pp. 1704–1731.

2. Khanal S, Ghimire P, Dhamoon AS. The Repertoire of Adenovirus in Human Disease: The Innocuous to the Deadly. Biomedicines. 2018; 6. doi:10.3390/biomedicines6010030

3. Chroboczek J, Bieber F, Jacrot B. The sequence of the genome of adenovirus type 5 and its comparison with the genome of adenovirus type 2. Virology. 1992;186: 280–285. doi:10.1016/0042-6822(92)90082-z

4. Davison AJ, Benko M, Harrach B. Genetic content and evolution of adenoviruses. J Gen Virol. 2003;84: 2895–2908. doi:10.1099/vir.0.19497-0

5. Berk AJ. Recent lessons in gene expression, cell cycle control, and cell biology from adenovirus. Oncogene. 2005;24: 7673–85. doi:10.1038/sj.onc.1209040

6. Chow LT, Gelinas RE, Broker TR, Roberts RJ. An amazing sequence arrangement at the 5’ ends of adenovirus 2 messenger RNA. Cell. 1977;12: 1–8.

7. Berget SM, Moore C, Sharp PA. Spliced segments at the 5’ terminus of adenovirus 2 late mRNA. PNAS. 1977;74: 3171–3175. doi:10.1073/pnas.74.8.3171

8. Sommer S, Salditt-Georgieff M, Bachenheimer S, Darnell JE, Furuichi Y, Morgan M, et al. The methylation of adenovirus-specific nuclear and cytoplasmic RNA. Nucleic Acids Res. 1976;3: 749–65.

9. Philipson L, Wall R, Glickman G, Darnell JE. Addition of polyadenylate sequences to virus-specific RNA during adenovirus replication. Proc Natl Acad Sci U S A. 1971;68: 2806–9.

10. Berk AJ. Discovery of RNA splicing and genes in pieces. Proc Natl Acad Sci USA. 2016;113: 801–805. doi:10.1073/pnas.1525084113

11. Montell C, Fisher EF, Caruthers MH, Berk AJ. Resolving the functions of overlapping viral genes by site-specific mutagenesis at a mRNA splice site. Nature. 1982;295: 380–384. doi:10.1038/295380a0

12. Winberg G, Shenk T. Dissection of overlapping functions within the adenovirus type 5 E1A gene. EMBO J. 1984;3: 1907–1912.

13. Fonseca GJ, Thillainadesan G, Yousef AF, Ablack JN, Mossman KL, Torchia J, et al. Adenovirus evasion of interferon-mediated innate immunity by direct antagonism of a cellular histone posttranslational modification. Cell Host Microbe. 2012; 11: 597–606. doi:10.1016/j.chom.2012.05.005

14. Zemke NR, Berk AJ. The Adenovirus E1A C Terminus Suppresses a Delayed Antiviral Response and Modulates RAS Signaling. Cell Host & Microbe. 2017;22: 789–800.e5. doi:10.1016/j.chom.2017.11.008

15. Pelka P, Ablack JNG, Fonseca GJ, Yousef AF, Mymryk JS. Intrinsic Structural Disorder in Adenovirus E1A: a Viral Molecular Hub Linking Multiple Diverse Processes. Journal of Virology. 2008;82: 7252–7263. doi:10.1128/JVI.00104-08

16. Blackford AN, Grand RJA. Adenovirus E1B 55-Kilodalton Protein: Multiple Roles in Viral Infection and Cell Transformation. Journal of Virology. 2009;83: 4000–4012. doi:10.1128/JVI.02417-08

17. Han J, Sabbatini P, Perez D, Rao L, Modha D, White E. The E1B 19K protein blocks apoptosis by interacting with and inhibiting the p53-inducible and death-promoting Bax protein. Genes Dev. 1996;10: 461–477. doi:10.1101/gad.10.4.461

18. Yew PR, Berk AJ. Inhibition of p53 transactivation required for transformation by adenovirus early 1B protein. Nature. 1992;357: 82–85. doi:10.1038/357082a0

19. Bridge E, Ketner G. Interaction of adenoviral E4 and E1b products in late gene expression. Virology. 1990;174: 345–53.

20. Cathomen T, Weitzman MD. A functional complex of adenovirus proteins E1B-55kDa and E4orf6 is necessary to modulate the expression level of p53 but not its transcriptional activity. J Virol. 2000;74: 11407–12.

21. Harada JN, Shevchenko A, Shevchenko A, Pallas DC, Berk AJ. Analysis of the adenovirus E1B-55K-anchored proteome reveals its link to ubiquitination machinery. J Virol. 2002;76: 9194–206.

22. Querido E, Blanchette P, Yan Q, Kamura T, Morrison M, Boivin D, et al. Degradation of p53 by adenovirus E4orf6 and E1B55K proteins occurs via a novel mechanism involving a Cullin-containing complex. Genes Dev. 2001;15: 3104–17. doi:10.1101/gad.926401

23. Dybas JM, Herrmann C, Weitzman MD. Ubiquitination at the interface of tumor viruses and DNA damage responses. Curr Opin Virol. 2018;32: 40–47. doi:10.1016/j.coviro.2018.08.017

24. Winnacker EL. Adenovirus DNA: structure and function of a novel replicon. Cell. 1978;14: 761–773. doi:10.1016/0092-8674(78)90332-x

25. Webster A, Leith IR, Nicholson J, Hounsell J, Hay RT. Role of preterminal protein processing in adenovirus replication. J Virol. 1997;71: 6381–6389.

26. Brenkman AB, Breure EC, van der Vliet PC. Molecular architecture of adenovirus DNA polymerase and location of the protein primer. J Virol. 2002;76: 8200–8207. doi:10.1128/jvi.76.16.8200-8207.2002

27. de Jong RN, van der Vliet PC, Brenkman AB. Adenovirus DNA replication: protein priming, jumping back and the role of the DNA binding protein DBP. Curr Top Microbiol Immunol. 2003;272: 187–211. doi:10.1007/978-3-662-05597-7_7

28. Robinson CM, Rajaiya J, Zhou X, Singh G, Dyer DW, Chodosh J. The E3 CR1-gamma gene in human adenoviruses associated with epidemic keratoconjunctivitis. Virus Res. 2011;160: 120–127. doi:10.1016/j.virusres.2011.05.022

29. Singh G, Robinson CM, Dehghan S, Jones MS, Dyer DW, Seto D, et al. Homologous recombination in E3 genes of human adenovirus species D. J Virol. 2013;87: 12481–12488. doi:10.1128/JVI.01927-13

30. Wold WSM, Tollefson AE, Hermiston TW. E3 Transcription Unit of Adenovirus. In: Doerfler W, Böhm P, editors. The Molecular Repertoire of Adenoviruses I: Virion Structure and Infection. Berlin, Heidelberg: Springer; 1995. pp. 237–274. doi:10.1007/978-3-642-79496-4_13

31. Bridge E, Ketner G. Redundant control of adenovirus late gene expression by early region 4. J Virol. 1989;63: 631–8.

32. Weitzman MD. Functions of the adenovirus E4 proteins and their impact on viral vectors. Front Biosci. 2005;10: 1106–17.

33. Weitzman MD, Ornelles DA. Inactivating intracellular antiviral responses during adenovirus infection. Oncogene. 2005;24: 7686–96. doi:10.1038/sj.onc.1209063

34. Weinmann R, Raskas HJ, Roeder RG. Role of DNA-dependent RNA polymerases II and III in transcription of the adenovirus genome late in productive infection. Proc Natl Acad Sci USA. 1974;71: 3426–3439. doi:10.1073/pnas.71.9.3426

35. Vachon VK, Conn GL. Adenovirus VA RNA: An essential pro-viral non-coding RNA. Virus Res. 2016;212: 39–52. doi:10.1016/j.virusres.2015.06.018

36. Hoeben RC, Uil TG. Adenovirus DNA Replication. Cold Spring Harb Perspect Biol. 2013;5. doi:10.1101/cshperspect.a013003

37. Shaw AR, Ziff EB. Transcripts from the adenovirus-2 major late promoter yield a single early family of 3’ coterminal mRNAs and five late families. Cell. 1980;22: 905–916. doi:10.1016/0092-8674(80)90568-1

38. Parks RJ. Adenovirus protein IX: a new look at an old protein. Mol Ther. 2005;11: 19–25. doi:10.1016/j.ymthe.2004.09.018

39. Zhang W, Imperiale MJ. Requirement of the adenovirus IVa2 protein for virus assembly. J Virol. 2003;77: 3586–3594. doi:10.1128/jvi.77.6.3586-3594.2003

40. Carter TH, Ginsberg HS. Viral transcription in KB cells infected by temperature-sensitive “early” mutants of adenovirus type 5. J Virol. 1976;18: 156–166.

41. Crossland LD, Raskas HJ. Identification of adenovirus genes that require template replication for expression. J Virol. 1983;46: 737–748.

42. Thomas GP, Mathews MB. DNA replication and the early to late transition in adenovirus infection. Cell. 1980;22: 523–533. doi:10.1016/0092-8674(80)90362-1

43. Ramke M, Lee JY, Dyer DW, Seto D, Rajaiya J, Chodosh J. The 5’UTR in human adenoviruses: leader diversity in late gene expression. Sci Rep. 2017;7: 618. doi:10.1038/s41598-017-00747-y

44. Soloway PD, Shenk T. The adenovirus type 5 i-leader open reading frame functions in cis to reduce the half-life of L1 mRNAs. J Virol. 1990;64: 551–558.

45. Morris SJ, Scott GE, Leppard KN. Adenovirus Late-Phase Infection Is Controlled by a Novel L4 Promoter. J Virol. 2010;84: 7096–7104. doi:10.1128/JVI.00107-10

46. Biasiotto R, Akusjärvi G. Regulation of Human Adenovirus Alternative RNA Splicing by the Adenoviral L4-33K and L4-22K Proteins. Int J Mol Sci. 2015;16: 2893–2912. doi:10.3390/ijms16022893

47. Tollefson AE, Ying B, Doronin K, Sidor PD, Wold WSM. Identification of a New Human Adenovirus Protein Encoded by a Novel Late l-Strand Transcription Unit. Journal of Virology. 2007;81: 12918–12926. doi:10.1128/JVI.01531-07

48. Ying B, Tollefson AE, Wold WSM. Identification of a Previously Unrecognized Promoter That Drives Expression of the UXP Transcription Unit in the Human Adenovirus Type 5 Genome. J Virol. 2010;84: 11470–11478. doi:10.1128/JVI.01338-10

49. Miller DL, Myers CL, Rickards B, Coller HA, Flint SJ. Adenovirus type 5 exerts genome-wide control over cellular programs governing proliferation, quiescence, and survival. Genome Biology. 2007;8: R58. doi:10.1186/gb-2007-8-4-r58

50. Miller DL, Rickards B, Mashiba M, Huang W, Flint SJ. The adenoviral E1B 55-kilodalton protein controls expression of immune response genes but not p53-dependent transcription. J Virol. 2009;83: 3591–3603. doi:10.1128/JVI.02269-08

51. Zhao H, Dahlö M, Isaksson A, Syvänen A-C, Pettersson U. The transcriptome of the adenovirus infected cell. Virology. 2012;424: 115–128. doi:10.1016/j.virol.2011.12.006

52. Zhao H, Chen M, Valdés A, Lind SB, Pettersson U. Transcriptomic and proteomic analyses reveal new insights into the regulation of immune pathways during adenovirus type 2 infection. BMC Microbiol. 2019;19: 15. doi:10.1186/s12866-018-1375-5

53. Zhao H, Chen M, Pettersson U. A new look at adenovirus splicing. Virology. 2014;456–457: 329–341. doi:10.1016/j.virol.2014.04.006

54. Crisostomo L, Soriano AM, Mendez M, Graves D, Pelka P. Temporal dynamics of adenovirus 5 gene expression in normal human cells. PLOS ONE. 2019;14: e0211192. doi:10.1371/journal.pone.0211192

55. Brandes N, Linial M. Gene overlapping and size constraints in the viral world. Biology Direct. 2016;11: 26. doi:10.1186/s13062-016-0128-3

56. O’Grady T, Wang X, Höner zu Bentrup K, Baddoo M, Concha M, Flemington EK. Global transcript structure resolution of high gene density genomes through multi-platform data integration. Nucleic Acids Res. 2016;44: e145–e145. doi:10.1093/nar/gkw629

57. Depledge DP, Srinivas KP, Sadaoka T, Bready D, Mori Y, Placantonakis DG, et al. Direct RNA sequencing on nanopore arrays redefines the transcriptional complexity of a viral pathogen. Nat Commun. 2019;10: 1–13. doi:10.1038/s41467-019-08734-9

58. Tombácz D, Balázs Z, Csabai Z, Moldován N, Szűcs A, Sharon D, et al. Characterization of the Dynamic Transcriptome of a Herpesvirus with Long-read Single Molecule Real-Time Sequencing. Sci Rep. 2017;7: 43751. doi:10.1038/srep43751

59. Viehweger A, Krautwurst S, Lamkiewicz K, Madhugiri R, Ziebuhr J, Hölzer M, et al. Direct RNA nanopore sequencing of full-length coronavirus genomes provides novel insights into structural variants and enables modification analysis. Genome Res. 2019;29: 1545–1554. doi:10.1101/gr.247064.118

60. Weirather JL, de Cesare M, Wang Y, Piazza P, Sebastiano V, Wang X-J, et al. Comprehensive comparison of Pacific Biosciences and Oxford Nanopore Technologies and their applications to transcriptome analysis. F1000Res. 2017;6: 100. doi:10.12688/f1000research.10571.2

61. Danecek P, Auton A, Abecasis G, Albers CA, Banks E, DePristo MA, et al. The variant call format and VCFtools. Bioinformatics. 2011;27: 2156–2158. doi:10.1093/bioinformatics/btr330

62. Narasimhan V, Danecek P, Scally A, Xue Y, Tyler-Smith C, Durbin R. BCFtools/RoH: a hidden Markov model approach for detecting autozygosity from next-generation sequencing data. Bioinformatics. 2016;32: 1749–1751. doi:10.1093/bioinformatics/btw044

63. Li X, Xiong X, Yi C. Epitranscriptome sequencing technologies: decoding RNA modifications. Nat Methods. 2017;14: 23–31. doi:10.1038/nmeth.4110

64. Garalde DR, Snell EA, Jachimowicz D, Sipos B, Lloyd JH, Bruce M, et al. Highly parallel direct RNA sequencing on an array of nanopores. Nat Methods. 2018;15: 201–206. doi:10.1038/nmeth.4577

65. Wu TD, Reeder J, Lawrence M, Becker G, Brauer MJ. GMAP and GSNAP for Genomic Sequence Alignment: Enhancements to Speed, Accuracy, and Functionality. Methods Mol Biol. 2016;1418: 283–334. doi:10.1007/978-1-4939-3578-9_15

66. Li H. Minimap2: pairwise alignment for nucleotide sequences. Bioinformatics. 2018;34: 3094–3100. doi:10.1093/bioinformatics/bty191

67. Bonfert T, Friedel CC. Prediction of Poly(A) Sites by Poly(A) Read Mapping. PLoS ONE. 2017;12: e0170914. doi:10.1371/journal.pone.0170914

68. Borodulina OR, Kramerov DA. Transcripts synthesized by RNA polymerase III can be polyadenylated in an AAUAAA-dependent manner. RNA. 2008;14: 1865–1873. doi:10.1261/rna.1006608

69. Tang AD, Soulette CM, Baren MJ van, Hart K, Hrabeta-Robinson E, Wu CJ, et al. Full-length transcript characterization of SF3B1 mutation in chronic lymphocytic leukemia reveals downregulation of retained introns. bioRxiv. 2018; 410183. doi:10.1101/410183

70. Hidalgo P, Anzures L, Hernández-Mendoza A, Guerrero A, Wood CD, Valdés M, et al. Morphological, Biochemical, and Functional Study of Viral Replication Compartments Isolated from Adenovirus-Infected Cells. Journal of Virology. 2016;90: 3411–3427. doi:10.1128/JVI.00033-16

71. Stern-Ginossar N, Weisburd B, Michalski A, Le VTK, Hein MY, Huang S-X, et al. Decoding human cytomegalovirus. Science. 2012;338: 1088–1093. doi:10.1126/science.1227919

72. Arvey A, Tempera I, Tsai K, Chen H-S, Tikhmyanova N, Klichinsky M, et al. An Atlas of the Epstein-Barr Virus Transcriptome and Epigenome Reveals Host-Virus Regulatory Interactions. Cell Host Microbe. 2012;12: 233–245. doi:10.1016/j.chom.2012.06.008

73. Arias C, Weisburd B, Stern-Ginossar N, Mercier A, Madrid AS, Bellare P, et al. KSHV 2.0: A Comprehensive Annotation of the Kaposi’s Sarcoma-Associated Herpesvirus Genome Using Next-Generation Sequencing Reveals Novel Genomic and Functional Features. PLOS Pathogens. 2014;10: e1003847. doi:10.1371/journal.ppat.1003847

74. Rutkowski AJ, Erhard F, L’Hernault A, Bonfert T, Schilhabel M, Crump C, et al. Widespread disruption of host transcription termination in HSV-1 infection. Nat Commun. 2015;6: 7126. doi:10.1038/ncomms8126

75. Garren SB, Kondaveeti Y, Duff MO, Carmichael GG. Global Analysis of Mouse Polyomavirus Infection Reveals Dynamic Regulation of Viral and Host Gene Expression and Promiscuous Viral RNA Editing. PLoS Pathog. 2015;11: e1005166. doi:10.1371/journal.ppat.1005166

76. Evans VC, Barker G, Heesom KJ, Fan J, Bessant C, Matthews DA. De novo derivation of proteomes from transcriptomes for transcript and protein identification. Nat Methods. 2012;9: 1207–1211. doi:10.1038/nmeth.2227

77. Wright J, Leppard KN. The Human Adenovirus 5 L4 Promoter Is Activated by Cellular Stress Response Protein p53. J Virol. 2013;87: 11617–11625. doi:10.1128/JVI.01924-13

78. Wright J, Atwan Z, Morris SJ, Leppard KN. The Human Adenovirus Type 5 L4 Promoter Is Negatively Regulated by TFII-I and L4-33K. Journal of Virology. 2015;89: 7053–7063. doi:10.1128/JVI.00683-15

79. Hennig T, Michalski M, Rutkowski AJ, Djakovic L, Whisnant AW, Friedl MS, et al. HSV-1-induced disruption of transcription termination resembles a cellular stress response but selectively increases chromatin accessibility downstream of genes. PLoS Pathog. 2018;14: e1006954. doi:10.1371/journal.ppat.1006954

80. Ramírez F, Ryan DP, Grüning B, Bhardwaj V, Kilpert F, Richter AS, et al. deepTools2: a next generation web server for deep-sequencing data analysis. Nucleic Acids Res. 2016;44: W160–165. doi:10.1093/nar/gkw257

81. Feng Y-Y, Ramu A, Cotto KC, Skidmore ZL, Kunisaki J, Conrad DF, et al. RegTools: Integrated analysis of genomic and transcriptomic data for discovery of splicing variants in cancer. bioRxiv. 2018; 436634. doi:10.1101/436634

82. Robinson JT, Thorvaldsdóttir H, Winckler W, Guttman M, Lander ES, Getz G, et al. Integrative Genomics Viewer. Nat Biotechnol. 2011;29: 24–26. doi:10.1038/nbt.1754

83. Li H, Handsaker B, Wysoker A, Fennell T, Ruan J, Homer N, et al. The Sequence Alignment/Map format and SAMtools. Bioinformatics. 2009;25: 2078–2079. doi:10.1093/bioinformatics/btp352

84. Quinlan AR, Hall IM. BEDTools: a flexible suite of utilities for comparing genomic features. Bioinformatics. 2010;26: 841–842. doi:10.1093/bioinformatics/btq033

85. Heinz S, Benner C, Spann N, Bertolino E, Lin YC, Laslo P, et al. Simple combinations of lineage-determining transcription factors prime cis-regulatory elements required for macrophage and B cell identities. Mol Cell. 2010;38: 576–589. doi:10.1016/j.molcel.2010.05.004

